# Novel *exc* Genes Involved in Formation of the Tubular Excretory Canals of *C. elegans*

**DOI:** 10.1101/359653

**Authors:** Hikmat Al-Hashimi, Travis Chiarelli, Erik A. Lundquist, Matthew Buechner

## Abstract

Regulation of luminal diameter is critical to the function of small single-celled tubes, of which the seamless tubular excretory canals of *C. elegans* provide a tractable genetic model. Mutations in several sets of genes exhibit the Exc phenotype, in which canal luminal growth is visibly altered. Here, a focused reverse genomic screen of genes highly expressed in the canals found 24 genes that significantly affect luminal outgrowth or diameter. These genes encode novel proteins as well as highly conserved proteins involved in processes including gene expression, cytoskeletal regulation, vesicular movement, and transmembrane transport. In addition, two genes act as suppressors on a pathway of conserved genes whose products mediate vesicle movement from early to recycling endosomes. The results provide new tools for understanding the integration of cytoplasmic structure and physiology in forming and maintaining the narrow diameter of single-cell tubules.

## INTRODUCTION

Tubule formation is an essential process during development of multicellular organisms, with the narrowest tubes occurring in structures as diverse as *Drosophila* trachea, floral pollen tubes, and mammalian capillaries (Lubarsky and Krasnow 2003; SigürbjÖrnsdÓttir *et al.* 2014). In *C. elegans*, the excretory system is comprised of cells that form single-celled tubules of three types: pore cells that wrap around a lumen to form a tube with an autocellular junction (“seamed tube”); a larger duct cell that forms a similar tube followed by dissolution of the junction to form a “seamless” tube; and the large excretory canal cell that extends four long seamless tubules (“canals”) throughout the length of the organism (Sundaram and Buechner 2016).

Many mutants have been discovered that affect the length, guidance of outgrowth, or lumen diameter of the excretory canals. An initial set of such identified “*exc*” mutants were mapped (Buechner *et al.* 1999), and found to include multiple alleles of some *exc* genes, but only single alleles of others. The frequency of mutations suggested that additional genes should have excretory lumen defects. Studies by multiple laboratories indeed found alleles of other genes with Exc phenotypes (Khan *et al.* 2013; Kolotuev *et al.* 2013; Armenti *et al.* 2014; Lant *et al.* 2015; Gill *et al.* 2016; Forman-Rubinsky *et al.* 2017). Almost all of the original *exc* genes have now been cloned (Suzuki *et al.* 2001; Berry *et al.* 2003; Fujita *et al.* 2003; Praitis *et al.* 2005; Tong and Buechner 2008; Mattingly and Buechner 2011; Shaye and Greenwald 2015; Grussendorf *et al.* 2016; Al-Hashimi *et al.* 2018), and found to affect multiple well-conserved cell processes, including cytoskeletal structures, ion channels, and vesicle recycling pathways. The initial screen sought primarily non-lethal genetic effects, but several of the subsequently identified genes were lethal when null.

RNAi studies have been particularly useful in determining roles of excretory canal genes where the null allele is lethal, such as the gene encoding the NHR-31 nuclear hormone receptor (Hahn-Windgassen and Van Gilst 2009), the ABI-1 Abelson-Interactor (McShea *et al.* 2013), and the PROS-1 transcription factor (Kolotuev *et al.* 2013). In addition, null mutations in genes that connect the excretory canal cell to the excretory duct (e.g. LET-4 (Mancuso *et al.* 2012) and LPR-1 (Forman-Rubinsky *et al.* 2017)) are lethal.

In order to identify other genes affecting the process of tubulogenesis and tubule maintenance in the excretory canals, we undertook a targeted genomic RNAi screen to identify excretory canal genes that exhibit lumen alterations (“Exc” phenotypes) when knocked down. This screen confirmed or identified 24 genes preferentially expressed in the canals that showed effects on lumen and/or outgrowth of the excretory canals, including 17 genes with no prior known phenotypic effects on the canals. In addition, two knockdowns suppressed effects of mutation of the *exc-5* vesicle-recycling gene, and therefore represent potential regulators of vesicle transport needed for single-cell tubulogenesis.

## MATERIALS AND METHODS

### Nematode genetics

*C. elegans* strains (Table 1) were grown by use of standard culture techniques on lawns of *Escherichia coli* strain BK16 (a streptomycin-resistant derivative of strain 0P50) on nematode growth medium (NGM) plates (Sulston and Hodgkin 1988). All strains were grown and evaluated for canal phenotypes at 20°C. Worms observed in this study were young adults or adults.

**Table 1.** List of strains used in this study, with genotype descriptions.

| STRAIN | GENOTYPE | DESCRIPTION | REFERENCE |
| --- | --- | --- | --- |
| BK36 | *unc-119(ed3)* III; *qpIs11[unc-119; P<sub>vha-1::gfp</sub>]* I | N2 with integrated GFP marker expressed in excretory canal cytoplasm | (MATTINGLY AND BUECHNER 2011) |
| BK540 | *rrf-3(pk1426)* II; *qpIs11 [unc-119; P<sub>vha-1::gfp</sub>]* I | RNAi-sensitized strain expressing GFP in canals | This study |
| BK541 | *sid-1(pk3321)* II; *qpIs11 [unc-119; P<sub>vha-1::gfp</sub>]* I | Systemic RNAi-impaired strain expressing GFP in canals | This study |
| BK542 | *exc-2(rh90)* X; *qpIs11 [unc-119; P<sub>vha-1::gfp</sub>]* I | *exc-2(rh90)* expressing GFP in canals | This study |
| BK543 | *exc-3(rh207)* X; *qpIs11 [unc-119; P<sub>vha-1::gfp</sub>]* I | *exc-3(rh207)* expressing GFP in canals | This study |
| BK544 | *exc-4(rh133)* *qpIs11 [unc-119; P<sub>vha-1::gfp</sub>]* I | *exc-4(rh133)* expressing GFP in canals | This study |
| BK545 | *exc-5(rh232)* IV; *qpIs11 [unc-119; P<sub>vha-1::gfp</sub>]* I | *exc-5(rh232)* expressing GFP in canals | This study |
| BK546 | *exc-7(rh252)* II; *qpIs11 [unc-119; P<sub>vha-1::gfp</sub>]* I | *exc-7(rh252)* expressing GFP in canals | This study |
| BK547 | BK540; *exc-2(rh90)* X | *exc-2(rh90)* expressing GFP in canals in RNAi-sensitized background | This study |
| BK548 | BK540; *exc-3(rh207)* X | *exc-3(rh207)* expressing GFP in canals in RNAi-sensitized background | This study |
| BK549 | BK540; *exc-4(rh133)* I | *exc-4(rh133)* expressing GFP in canals in RNAi-sensitized background | This study |
| BK550 | BK540; *exc-5(rh232)* IV | *exc-5(rh232)* expressing GFP in canals in RNAi-sensitized background | This study |

Each nematode strain (wild-type N2, and *exc-2, exc-3, exc-4, exc-5, and exc-7*) was crossed to strain BK36, which harbors a chromosomal insertion of a canal-specific promoter driving cytoplasmic GFP expression (P*vha-1::gfp).* Strains were then sensitized for RNAi treatment by crossing them to mutant strain BK540 (a strain carrying *rrf-3*(*pk1426*) in addition to the same chromosomal *gfp* insertion as above) and selecting in the F2 generation for homozygous *rrf-3* deletion allele and appropriate *exc* mutation. (As *exc-7* maps very close to *rrf-3*, the *exc-7* strain carrying *gfp* was not crossed to BK540 and was not sensitized to RNAi). For all sensitized strains, the *rrf-3* deletion was confirmed via PCR using the forward primer ^5’^GCTGGATATTGCCGAGCAC^3’^, reverse primer ^5’^GGAGATCTCCGAGCCCTAGAC^3^’, and a reverse nested primer ^5’,^CATCGCCAGGCCAACTCAATAC^3’^. As a negative control, we crossed BK36 to RNAi-refractive strain NL3321 *sid-1*(*pk3321*).

### RNAi Screen

The Ahringer RNAi bacterial library (Kamath *et al.* 2003) was utilized for this study. Overnight cultures were prepared by inoculating bacteria in 5 ml LB + ampicillin (100μg/ml) + tetracycline (12.5μg/ml), and cultured at 37°C for 16 hours. In order to induce the bacteria with IPTG, overnight cultures were moved to fresh media, incubated at 37°C with rotation until cultures reached an O.D._600_ in a range from 0.5 to 0.8. IPTG was then added to the culture to a final concentration of 95 μg/ml along with ampicillin at 100μg/ml. The cultures were then incubated with rotation at 37°C for ninety minutes followed by re-induction with IPTG and ampicillin, and another ninety minutes of incubation at 37°C with rotation. Finally, IPTG and ampicillin were added for the last time right before using these bacteria to seed NGM in 12-well plates and Petri dishes. Plates were then incubated at room temperature for 24 hours in order to dry. L2 worms were added to the plates, and their F1 progeny were evaluated for phenotypes in the excretory canals. Each set of genes tested was induced together with induction of the *sid-1* negative control strain BK541 and of two positive control strains: a plate of bacteria induced to knock down *dpy-11* (which affects the hypoderm but not the canals) (Brenner 1974), and a plate of bacteria induced to knock down *erm-1* bacteria, which causes severe defects in excretory canal length and lumen diameter (Khan *et al.* 2013), respectively. Induction was considered successful and plates were screened only if worms grown on the control plates showed the appropriate phenotypes in at least 80% of the surviving progeny.

For each tested gene, the induced bacteria were seeded on one 12-well plate and one 60mm plate. Two or three L2 nematodes were placed on the bacterial lawn of each well, and screened for phenotypes in the 4^th^, 5^th^, and 6^th^ days of induction. Each gene was tested via RNAi treatment of twelve different strains of worms, shown in Table 1, while the sole 60mm plate was used for further analysis of animals with wild-type canals (strain BK540, Table 1) grown on the RNAi-expressing bacteria. For assessment of a canal effect, a minimum of five animals showing a canal phenotype were collected, examined, and in most cases photographed; for most genes, 10-20 affected animals were examined closely. For the 24 genes showing effects, the entire experiment was subsequently repeated, with induction and growth of bacteria solely on 60mm plates and feeding tested on BK540 (RNAi-sensitized wild-type with integrated canal marker) worms.

### Microscopy

Living worms were mounted on 3% agarose pads to which were added 0.1μ,m-diameter Polybead^®^ polystyrene beads (Polysciences, Warrington, PA) to immobilize the animals (Kim *et al.* 2013). Images were captured with a MagnaFire Camera (Optronics) on a Zeiss Axioskop microscope equipped with Nomarski optics and fluorescence set to 488 nm excitation and 520 nm emission. Adobe Photoshop software was used to combine images from multiple sections of individual worms and to crop them. Contrast on images was uniformly increased to show the excretory canal tissue more clearly.

### Canal Measurements

For measuring effects of suppression of the Exc-5 phenotype, excretory canal length and cystic and suppression phenotypes were measured and analyzed as described (Tong and Buechner 2008). Canal length was scored by eye on a scale from 0-4: A score of (4) was given if the canals had grown out to full length; canals that extended halfway past the vulva (midbody) to full-length were scored as (3); at the vulva (2); canals that ended halfway between the cell body and the vulva were scored as (1); and if the canal did not extend past the cell body, the canal was scored as (0). For statistical analyses, canals were binned into three categories for length (scores 0-1, scores 1.5-3.0, and score 3.5-4), and the results then analyzed via a 3×2 Fisher’s Exact Test (www.vassarstats.net).

### Reagent and Data Availability

All nematode strains used in this study are listed in Table 1. Bacterial clone numbers tested, and summary of test results are presented in Tables S1 and S2, available on Figshare. Gene names *exc-10* through *exc-18* and *suex-1* and *suex-2* have been registered with Wormbase (www.wormbase.org). Sensitized *exc* mutant strains are available upon request, and may be made available through the Caenorhabditis Genetics Center (CGC), University of Minnesota (cgc.umn.edu), pending acceptance to that repository. Other strains are available upon request.

## RESULTS AND DISCUSSION

### A focused RNAi screen for new *exc* mutations

A study of genomic expression in *C. elegans* was previously undertaken by the Miller lab (Spencer *et al.* 2011). In that study, lists of genes highly expressed in various tissues, including 250 genes preferentially expressed in the excretory canal cell, were made public on the website WormViz (http://www.vanderbilt.edu/wormdoc/wormmap/WormViz.html). Of the corresponding strains in the Ahringer library of bacteria expressing dsRNA to specific *C. elegans* genes (Kamath *et al.* 2003), we found that 216 grew well, and were tested for effects on the various *C. elegans* strains (Table S1).

The excretory canal cell has some characteristics similar to those of neurons: long processes guided by netrins and other neural guidance cues (Hedgecock *et al.* 1987), as well as early expression of the gene EXC-7/HuR/ELAV (Fujita *et al.* 2003), and so was considered potentially refractory to feeding RNAi (Calixto *et al.* 2010). We crossed strain BK36, containing a strong canal-specific integrated *gfp* marker, to a mutant in the *rrf-3* gene (*pk1426*) in order to increase sensitivity to RNAi (Simmer *et al.* 2002) to create strain BK540. In addition, we also crossed the same *gfp* marker and *rrf-3* mutation to excretory canal mutants *exc-2, exc-3, exc-4, exc-5*, and *exc-7* (except that *exc-7* was not RNAi-sensitized; see Materials and Methods). This was done in order to determine if the tested gene knockdowns interacted with known *exc* genes affecting excretory canal tubulogenesis, since double mutants in some *exc* genes (e.g. *exc-3; exc-7* double mutants (Buechner *et al.* 1999)) exhibit more severe canal phenotypes than either mutant alone.

We demonstrated the effectiveness of the treatment by performing successful knockdowns of canal-specific and -non-specific genes in these strains. Control knockdowns of *dpy-11* resulted in short worms with normal canal phenotypes, while knockdown of *exc-1* caused formation of variable-sized cysts in a shortened excretory canal, with no other obvious phenotypes (Fig. 1). Here, knockdown of the ezrin-moesin-radixin homologue gene *erm-1* (Göbel *et al.* 2004; Khan *et al.* 2013) caused severe malformation of the canals visible in 80% of surviving treated worms. A deletion mutant of this gene is often lethal due to cystic malformation of the intestine as well as the canals (Göbel *et al.* 2004), while our treatment allowed many animals to survive to adulthood and reproduce. This result is consistent with our RNAi treatment causing variable levels of gene knockdown (Timmons and Fire 1998) in the excretory canals.

**Figure 1.**
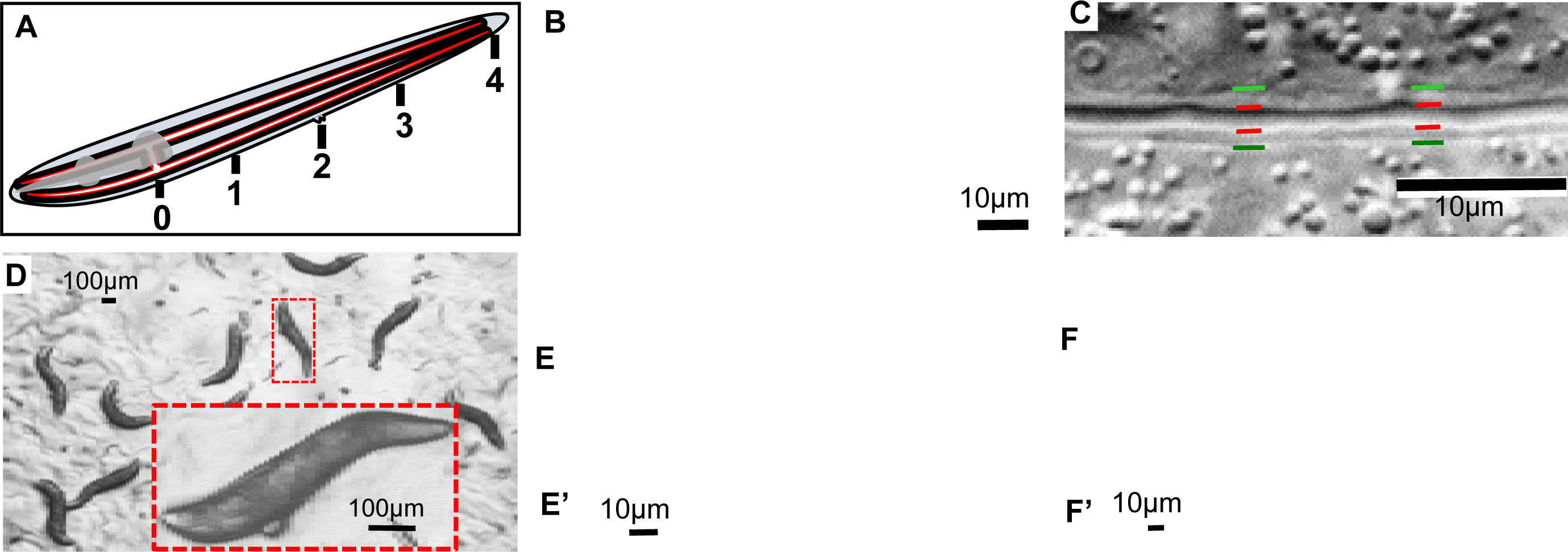
The excretory canals and induction controls. (A) Schematic diagram of the excretory canals extending over the full length of the worm with basal membrane (black) and apical membrane (red) surrounding a narrow lumen (white). Numbers 0-4 represent numerical assignments used to assess canal length. (B) DIC image of section of posterior excretory canal of wild-type worm (N2); canal is narrow with uniform diameter. Bar, 10μm. (C) Magnified DIC image of excretory canal of wild-type worm (N2). Lines indicate boundaries of canal lumen/apical surface (red) and cytoplasmic/basal surface (green). (D-F) Controls to ensure strong induction of dsRNA synthesis for RNAi screen, in *rrf-3(pk1426)* animals expressing GFP in the canals: (D) Knockdown of cuticle collagen gene *dpy-11.* Boxed image: Magnification of single worm. Bar = 100μm. (E) DIC and (E’) GFP image of *erm-1* knockdown. Bar, 10μm. (F) DIC and (F’) GFP image of *exc-1* knockdown. Bar, 10μm.

Of the 212 non-control genes tested, 182 caused no obvious phenotypic changes to the canals of BK540 worms, and 4 gave very low numbers (less than 5) of animals with mild defects. Knockdown of 24 genes caused noticeable defects in the development of the excretory canals in at least 5-10 worms, and gave this result upon retesting of these strains (Table 2, Table S2). The length of the canals was rated according to a measure shown in Fig. 1A, in which no extension past the excretory cell body was rated 0, extension to the animal midbody marked by the position of the vulva was measured as 2, and full extension was rated as 4. The average canal length of affected animals was characteristic for the gene knocked down (Table 2), although RNAi knockdown via feeding is intrinsically variable in the strength of gene induction and amount of bacteria eaten, so the observed canal length is likely longer than if the gene were fully and uniformly knocked out. Diameter of the canals also varied greatly, depending on the gene knocked down; in cases where fluid-filled cysts became evident (as in previously-described *exc* mutants), cyst size was rated as large (cyst diameter at least half the width of the animal), medium (one-quarter to one-half animal width), or small (up to one-quarter animal width).

**Table 2.** Genes tested exhibiting excretory canal defects via RNAi feeding ***. Genes previously found to have effects on excretory canal function or development are marked with an asterisk.

| Gene | Clone | Protein Class<br>Short Description of Known or<br>Inferred Protein Function | Canal RNAi Phenotype | # canals<br>examined | %<br>mutant<br>canals | Ave.<br>mutant<br>canal<br>length |
| --- | --- | --- | --- | --- | --- | --- |
| **Transcriptional and post-transcriptional factors** |  |  |  |  |  |  |
| *ceh-6\** | K02B12.1 | homeobox transcription factor | large fluid-filled cysts | Control |  |  |
| *ceh-37\** | C37E2.5 | Otx homeobox transcription factor | periodic cytoplasmic beads | 40 | 30% | 3.3 |
| *fbxa-183* | F44E7.6 | F-box protein, possible effects on RNA | swollen tip with vesicles | 125 | 66% | 3.1 |
| *mxt-1* | Y18D10A.8 | translation regulation | cytoplasmic beads with vesicles | 59 | 93% | 2.6 |
| **Cytoskeletal proteins and regulators** |  |  |  |  |  |  |
| *egal-1* | C10G6.1 | Egalitarian exonuclease, regulates dynein | medium-sized fluid-filled cysts | 40 | 100% | 1.1 |
| *cyk-1\** | F11H8.4 | *Diaphanous* formin | vesicles along swollen cytoplasm | 77 | 77% | 3.1 |
| **Transporters, channels, and receptors** |  |  |  |  |  |  |
| *inx-12\** | ZK770.3 | innexin gap junction protein | periodic cytoplasmic beads | 46 | 96% | 2.7 |
| *inx-13\** | Y8G1A.2 | innexin gap junction protein | periodic cytoplasmic beads | 50 | 80% | 2.8 |
| *vha-5\** | F35H10.4 | vacuolar ATPase component | beads, vesicles, swollen cytoplasm | 78 | 59% | 2.5 |
| *best-3* | C01B12.3 | bestrophin chloride channel | Swollen luminal tip | 36 | 28% | 3.4 |
| *exc-11* | F41E7.1 | Na<sup>+</sup>/H<sup>+</sup> solute carrier (SLC9 family) | medium-sized fluid-filled cysts | 34 | 97% | 1.7 |
| *exc-13* | T19D12.9 | sialic acid solute carrier (SLC17 family) | swollen tip, vesicles, convolutions | 23 | 22% | 3.3 |
| *exc-18* | C09F12.3 | Nematode 7tm GPCR | vesicles along swollen cytoplasm | 40 | 13% | 3.1 |
| **Vesicle movement regulators** |  |  |  |  |  |  |
| *gck-3\** | Y59A8B.23 | germinal center kinase protein | swollen tip, vesicles, convolutions | 76 | 57% | 3.5 |
| *mop-25.2* | Y53C12A.4 | scaffolding for endocytic recycling | medium-sized fluid-filled cysts,<br>vesicles along swollen cytoplasm | 20 | 100% | 1.1 |
| **Enzymatic activities** |  |  |  |  |  |  |
| *dhhc-2* | Y47H9C.2 | Zn-finger, palmitoyltransferase | cytoplasmic beads with vesicles | 57 | 21% | 3.5 |
| *gsr-1* | C46F11.2 | glutathione reductase | swollen tip, vesicles, convolutions | 73 | 5% | 3.4 |
| *gst-28* | Y53F4B.31 | glutathione-S-transferase, prostaglandin isomerase | swollen tip with vesicles | 26 | 27% | 3.4 |
| *exc-10* | T25C8.1 | sedoheptulose kinase | large fluid-filled cysts | 63 | 68% | 3.3 |
| *exc-14* | K11D12.9 | RING finger, possible E3 ubiquitin ligase | vesicles along swollen cytoplasm | 36 | 100% | 1.4 |
| *exc-15* | T08H10.1 | aldo-keto reductase | vesicles along swollen cytoplasm | 27 | 15% | 3.4 |
| Unknown function |  |  |  |  |  |  |
| *exc-12* | T05D4.3 | Nematode-only transmembrane protein | medium-sized fluid-filled cysts | 53 | 79% | 2.6 |
| *exc-16* | H09G03.1 | *Caenorhabditis*-only protein | vesicles along swollen cytoplasm | 134 | 7% | 3.3 |
| *exc-17* | C03G6.5 | Nematode conserved-domain protein | vesicles along swollen cytoplasm | 54 | 20% | 3.5 |
| Suppressors of *exc-5* mutation |  |  |  |  |  |  |
| *suex-1* | F12A10.7 | unknown *Caenorhabditis* protein | suppresses *exc-5* mutant cysts |  |  |  |
| *suex-2* | C53B4.1 | solute carrier (SLC22 family) | suppresses *exc-5* mutant cysts |  |  |  |

Finally, use of feeding RNAi knockdown allowed observation of gene effects where knockouts have been reported to be lethal.

### Excretory Canal Phenotypes

The common feature of all of these knockdown animals is that the posterior canals did not extend fully to the back of the animal (Table 2). The length of the canal lumen was often the same as the length of the canal cytoplasm, but in many cases the visible lumen (seen as a dark area in the center of the GFP-labelled cytoplasm) was substantially shorter than the length of the canal cytoplasm.

In addition to effects on canal length, the shape and width of the canal lumen and/or canal cytoplasm was affected by specific gene knockdown: A) Several knockdowns resulted in the formation of fluid-filled cysts reminiscent of those in known *exc* mutants; B) canals appeared normal in diameter, but had frequent thickenings of cytoplasm around otherwise normal (but short) lumen similar to the ‘beads” or “pearls” seen in growing first-larval-stage canals or in canals of animals undergoing osmotic stress (Kolotuev *et al.* 2013); C) Canal lumen ending in a large swelling of convoluted tubule or a multitude of small vesicles, and often with a “tail” of very thin canal cytoplasm without any lumen continuing distally; D) a series of vesicles filling much of the cytoplasm outside the normal-diameter lumen, and; E) an irregular shape of the basal surface of the cytoplasm, varying widely in diameter. Each of these phenotypes will be discussed below, together with the genes whose knockdown resulted in that phenotype.

CYSTIC CANALS: Two gene knockdowns, of *ceh-6* and of T25C8.1 (which will be referred to as *exc-10*) resulted in the formation of large fluid-filled cysts (Fig. 2), similar to those seen in *exc-2, exc-4*, and *exc-9* mutants (encoding an intermediate filament, a CLC chloride channel, and a CRIP vesicle-trafficking protein, respectively (Tong and Buechner 2008; Berry *et al.* 2003, Al-Hashimi, in press). The homeobox gene *ceh-6* encodes a well-studied transcription factor that defines expression of many genes in the canal (Burglin and Ruvkun 2001; Armstrong and Chamberlin 2010). Null mutants of this gene are lethal. The knockdowns had very short canals with large fluid-filled cysts. The effect of *ceh-6* knockdown could reflect lower transcription of many of the known *exc* genes, effects on the excretory aquaporin *aqp-8* (Mah *et al.* 2007), or of a novel gene.

**Figure 2.**
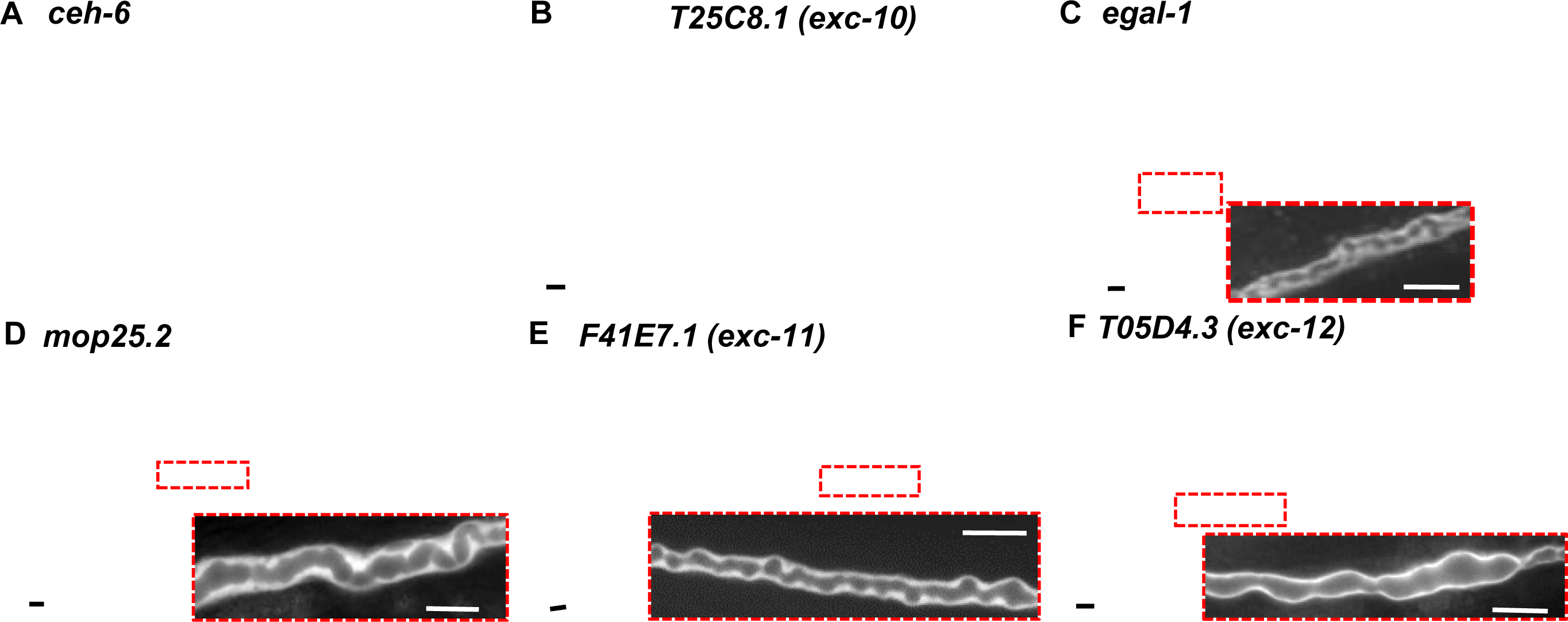
RNAi knockdowns causing formation of fluid-filled cysts or swollen lumen. (A-F) DIC images and (A’-F’) GFP fluorescence of representative animals exhibiting RNAi knockdown phenotypes: (A) *ceh-6*; (B) *T25C8.1 (exc-10);* (C) *egal-1;* (D) *mop-25.2;* (E) *F41E7.1 (exc-11);* (F) *T05D4.3 (exc-12).* Arrows: Medium and large fluid-filled cysts. All bars, 10 μm.

The second gene, T25C8.1 *(exc-10)* encodes a carbohydrate kinase (homology to sedoheptulose kinase) of unknown function in nematodes, although the human homologue SHPK has been linked to a lysosomal storage disease (Phornphutkul *et al.* 2001; Wamelink *et al.* 2008).

Knockdowns in *mop-25.2, egal-1*, F41E7.1 (*exc-11*), and T05D4.3 (*exc-12*) exhibited small-to-medium sized cysts (Fig. 2). In these animals, cystic regions of the lumen often appear to contain a series of hollow spheres, which may be connected or separate from each other along the lumen length (Fig. 2C-F). The EGAL-1 protein is a homologue of the *Drosophila* Egalitarian exonuclease involved in RNA degradation. EGAL-1 also interacts with dynein as part of a dynein-regulating complex at the face of the nucleus (Fridolfsson *et al.* 2010) and regulates polarity of the *Drosophila* egg chamber through organization of oocyte microtubules (Sanghavi *et al.* 2016). The excretory canal cell is rich in microtubules along the length of the canals (Buechner *et al.* 1999; Shaye and Greenwald 2015).

MOP-25.2 is a protein with close homology to yeast Mo25 and its homologues in all animals, and acts as a scaffolding protein for activating kinases including germinal center kinase at the STRIPAK complex, which also regulates RAB-11-mediated endocytic recycling in the excretory canal morphology and gonadal lumen formation in *C. elegans* (Lant *et al.* 2015; Pal *et al.* 2017). The *Drosophila* Mo25 also regulates transepithelial ion flux in the osmoregulatory Malpighian tubules (Sun *et al.* 2018).

F41E7.1 (*exc-11*) encodes a solute carrier with high homology to the Na^+^/H^+^ exchanger. The excretory cell lumen is associated with small canaliculi that have high levels of the vacuolar ATPase to pump protons into the canal lumen (Oka *et al.* 2001), so the presence of a Na^+^/H^+^ exchanger could be used for canal osmoregulatory function as well as luminal shape.

Finally, T05D4.3 (*exc-12*) is homologous only to genes in other nematodes, and has no obvious function, other than the presence of several putative transmembrane domains.

PERIODIC CYTOPLASMIC SWELLINGS: These “beads” or “pearls” are commonly seen in wild-type animals with growing canals at the L1 stage and in animals under osmotic stress (Kolotuev *et al.* 2013). These sites are hypothesized to be locations of addition of membrane to allow the canal to continue to grow together with the animal. The knockdown animals here were measured in young adulthood, and so should not exhibit such beads. Knockdown of the *inx-12* or *inx-13* genes (Fig. 3), which encode innexins highly expressed in the canals (and in the adjacent CAN neurons), gave rise to these structures. Innexins form the gap junctions of invertebrates (Hall 2017), and the excretory canals are rich in these proteins along the basal surface, where they connect the canal cytoplasm to the overlaying hypodermis (Nelson *et al.* 1983). Null mutants in either of these two genes results in early larval rod-like swollen lethality consistent with excretory cell malfunction (Altun *et al.* 2009). The knockdown phenotype here further suggests that these proteins regulate balancing of ionic content to allow normal canal growth.

**Figure 3.**
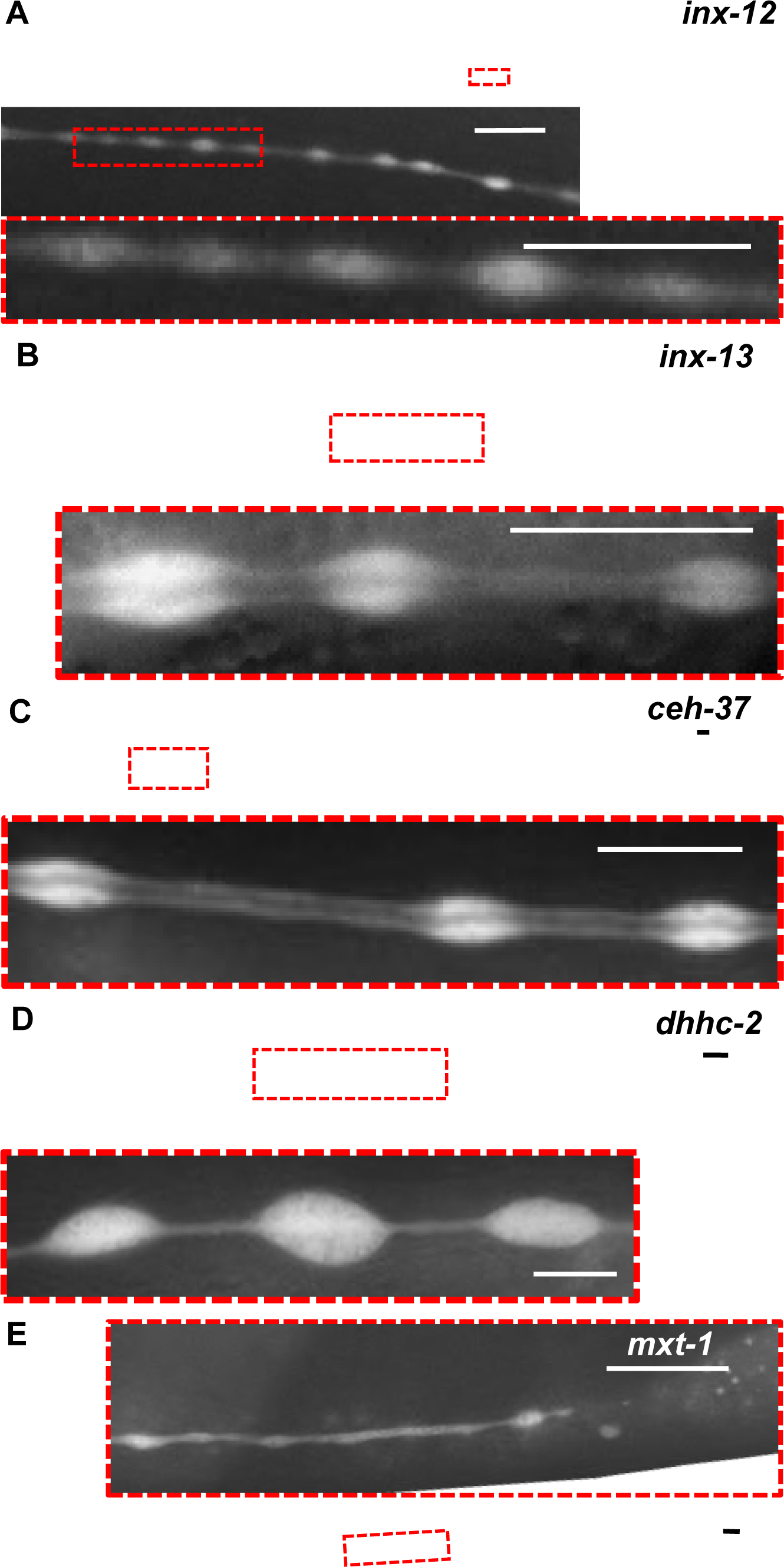
RNAi knockdowns causing periodic cytoplasmic swellings. GFP fluorescence images of swellings (“beads”) along length of canals. Boxed insets of marked areas are magnified to show width of lumen in regions within and between beads:. (A) *inx-12*; *inx-13;* (C) *ceh-37;* (D) *dhhc-2*; (E) *mxt-1.* Inset in (D) is of region posterior to end of lumen, so lumen is not visible. Note visible vesicles in cytoplasmic beads in (D). All bars, 10 μm.

A similar phenotype is seen in animals knocked down for *ceh-37*, which encodes a well-conserved otx Homeobox protein expressed solely in the excretory cell in adults, but additionally in a wide range of tissues in embryos (Lanjuin *et al.* 2003; Hench *et al.* 2015), and which binds preferentially to telomeric DNA (Moon *et al.* 2014). Expression of *ceh-37* is itself regulated by CEH-6 (Burglin and Ruvkun 2001), so the difference in phenotypes of knockdowns of these two genes suggests that CEH-6 regulates a wider range of genes than does CEH-37.

Knockdown of *dhhc-2* shows a similar phenotype, although bead size and placement appears more irregular than for the above knockdowns (Fig. 3). Close examination of the beads shows numerous small dark spots consistent with the presence of many vesicles of varying sizes within the beads (Fig. 3D) This gene encodes a zinc-finger protein homologous orthologous to human ZDHHC18, which acts as a protein palmitoyltransferase, possibly for small GTPase proteins (Ohno *et al.* 2012). A previous knockdown study of this gene (Edmonds and Morgan 2014) showed no obvious effects on morphology or behaviour, although combined knockdown of both *dhhc-2* together with its close homologue *dhhc-8* resulted in reduced lifespan for unknown reasons.

Finally, the “bead” phenotype is also seen in knockdown of *mxt-1*, an RNA-binding protein that binds to eukaryotic initiation factor 4E to regulate translation rates (Peter *et al.* 2015) (Fig. 3E).

SWELLING AT END OF LUMEN: The largest group of knockdown animals showed a substantial swelling at the distal tip of generally normal-diameter canals (Fig. 4). In some cases, the swelling appears to be caused by accumulation of a convoluted lumen folded back on itself, while in other knockdowns this swelling could reflect accumulation of a large number of vesicles at the end of the lumen. A combination of these structures also appears in many animals. Reflecting the variable effects of RNAi knockdown, some animals knocked down in genes discussed above, *ceh-37* and *mop25.2*, sometimes showed a highly convoluted lumen primarily at the distal tip (Fig. 4C, 4H), possibly reflecting weaker knockdown than in other examples where the entire lumen was affected. Knockdown of another gene, *best-3*, showed a similar effect (Fig. 4G). This gene encodes one of a large family of chloride channels homologous to human bestrophins, chloride channels found in muscle, neurons, and the eye, that are essential for Ca^++^ signaling, and defective in retinal diseases (Strauss *et al.* 2014).

**Figure 4.**
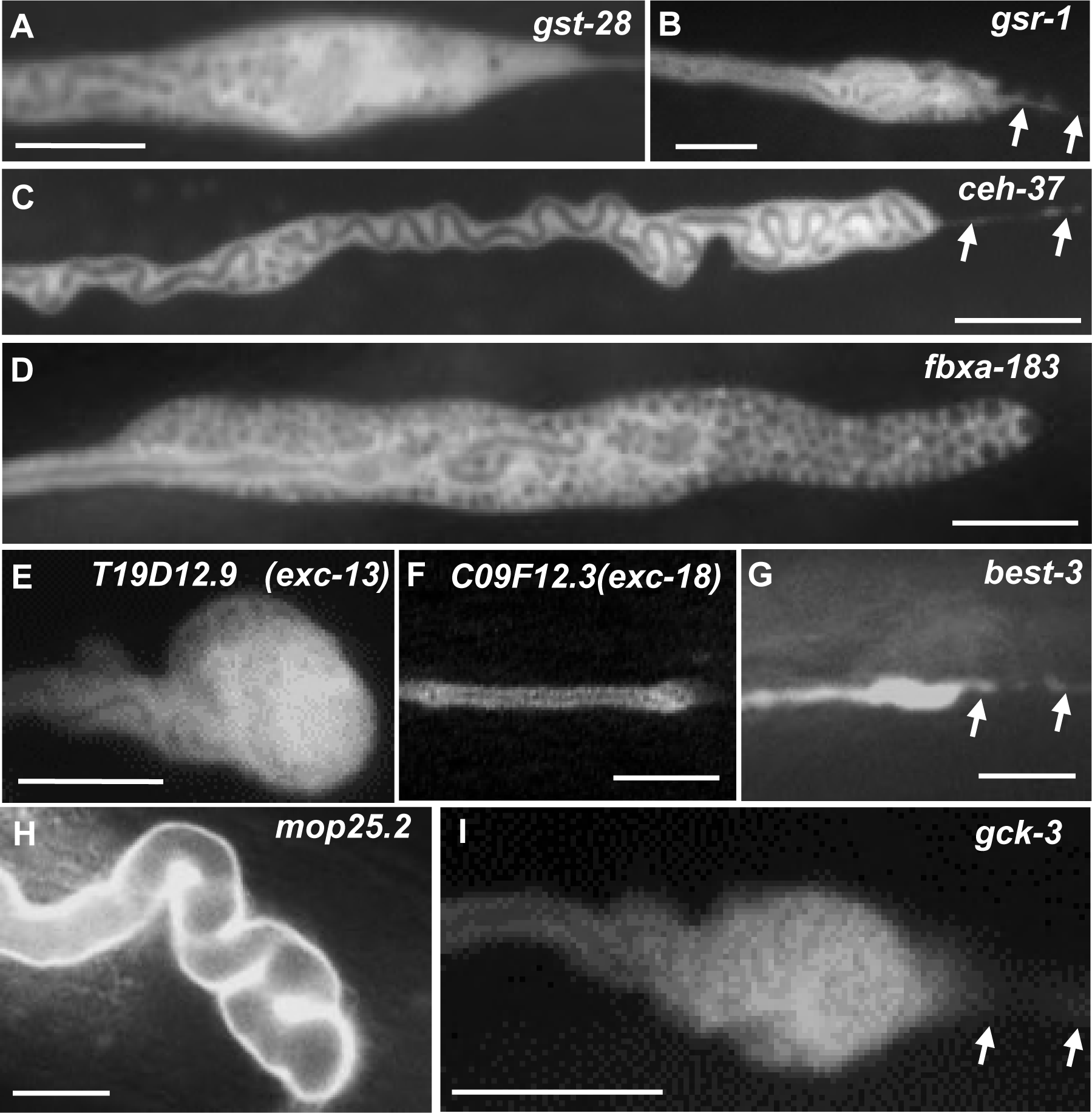
RNAi knockdowns causing swelling at end of lumen. GFP fluorescence images of swollen canals at termination of lumen caused by RNAi knockdown of genes: (A) *gst-28;* (B) *gsr-1;* (C) *ceh-37;* (D) *fbxa-183;* (E) *T19D12.9 (exc-13);* (F) *C09F12.3* (*exc-18*); (G) *best-3*; (H) *mop-235.2*; (I) *gck-3.* All images show regions of convoluted canals. Some areas in panels B, E, F, and I show additional areas that are appear as individual separated small cysts or large vesicles. Arrows: Cytoplasmic tail continuing past termination of lumen in panels B, C, G, and I. All bars, 10 μm.

Knockdowns of *gst-28* and of *fbxa-183* (Fig. 4A, 4D) show clear and dramatic accumulation of vesicles at the swelling at the tip of the lumen. Vesicle transport defects are the cause of canal malformations in *exc-1, exc-5*, and *exc-9* mutants (Tong and Buechner 2008; Mattingly and Buechner 2011; Grussendorf *et al.* 2016), so the knockdown effects shown in these and the following genes may reflect similar defects in vesicular transport. GST-28 is a glutathione-S-transferase orthologous to the human prostaglandin D synthase, which isomerizes PGH2 to form prostaglandin (Chang *et al.* 1987). FBXA-183 is one of the very large family of F-box proteins in *C. elegans* that facilitate targeting substrates for E3 ubiquitinase-mediated destruction (Kipreos and Pagano 2000). FBXA-183 also contains an FTH (FOG-2 Homology) domain; in FOG-2, this domain binds GLD-1, an RNA-binding protein (Clifford *et al.* 2000).

Knockdown of three genes produced animals with swollen distal canal tips filled with a mixture of convoluted tubule and individual vesicles. The *gsr-1* gene (Fig. 4B) encodes the sole glutathione reductase in *C. elegans*, necessary for rapid growth and embryonic development as well as canal morphology (Mora-Lorca *et al.* 2016). T19D12.9 (to be referred to as *exc-13*) (Fig. 4E) encodes another homologue of the human SLC family of solute carriers, including the ubiquitous lysosomal membrane sialic acid transport protein sialin (SLC17A5), which transports sialic acid from the lysosome, and nitrate from the plasma membrane in humans (Qin *et al.* 2012). C09F12.3 (to be referred to as *exc-18*) encodes a protein found predominantly in nematodes and likely encoding a 7-tm G-Protein-Coupled Receptor of the FMRFamide class, used to respond to the wide range of FMRFamide-Like Peptides (FLP) mediating multiple behaviors in invertebrates (Peymen *et al.* 2014). Finally, knockdown of the germinal center kinase gene *gck-3* (Fig. 4I) caused this phenotype. As noted above, germinal center kinase acts at the STRIPAK complex to regulate maintain canal morphology, and malformations affecting STRIPAK cause tubule defects such as cavernous cerebral malformations in humans (Lant *et al.* 2015).

A very narrow canal “tail” completely lacking a visible lumen often extends substantially past the end of the lumenated portion of the canal in these animals (Fig. 4B, 4C, 4G, 4I). This tail follows the path of wild-type canal growth, and in a few rare instances even reaches the normal endpoint of the canal. In wild-type animals, the lumen and tip of the canal grow together and reach the same endpoint (Buechner *et al.* 1999), with a widening suggestive of a growth cone at the tip of the growing canal in the embryo and L1 stage (Fujita *et al.* 2003). The tip of the canal is enriched in the formin EXC-6, which mediates interactions between microtubules and actin filaments and may mediate connections between the canal tip and end of the lumen (Shaye and Greenwald 2015). The results here are consistent with the idea that canal lumens grow and extend separately from the growing basal surface that guides cytoplasmic outgrowth (Kolotuev *et al.* 2013).

A NOVEL EXCRETORY PHENOTYPE: VESICLES ALONG LENGTH OF SWOLLEN CANAL CYTOPLASM: Knockdown of some genes gave rise to a phenotype that has not, to our knowledge, been observed before within the excretory canals (Fig. 5). Knockdown of the gene K11D12.9 (which will be referred to as *exc-14*) exhibited an extraordinary increase of vesicles in the cytoplasm of the canal, terminating in a large irregular swelling at the end of the canal (Fig. 5A, 5A’). This swelling is unusual in that the lumen of the canal appears relatively normal in diameter (though short), but is surrounded by cytoplasm that puffs out at the basal side of the cell, which is surrounded by (and extensively connected via innexins to) the hypoderm and by basement membrane abutting the pseudocoelom (Nelson *et al.* 1983). GFP labelling of the cytoplasm showed the thick layer of canaliculi surrounding the lumen, which is surrounded by a cytoplasm packed with vesicles of variable size. K11D12.9 encodes a protein containing a RING finger domain at the C-terminus, with BLASTP homology to potential ubiquitin E3-ligases found in plants and animals.

**Figure 5.**
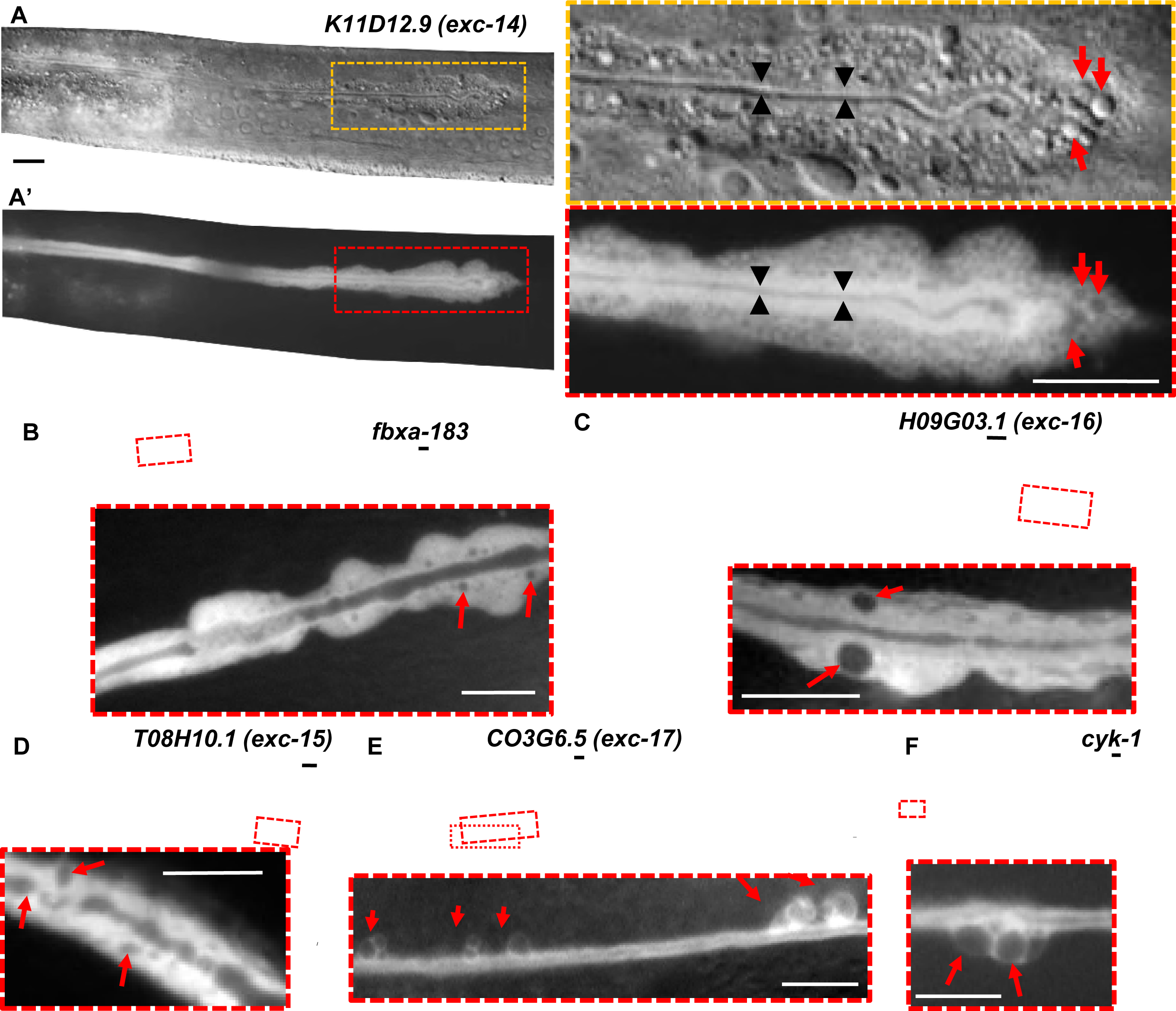
RNAi knockdowns causing irregular basal membrane along canal length. (A) DIC and (A’) GFP fluorescence images of distal tip of canal of representative animal knocked down for *K11D12.9 (exc-14).* Boxed areas are enlarged to right. Thin lumen indicated by black arrowheads is surrounded by area of bright GFP fluorescence. Distorted cytoplasmic shape is filled with large number of vesicles (red arrows). (B-F) GFP fluorescence of representative animals knocked down for genes: (B) *fbxa-183;* (C) *H09G03.1(exc-16);* (D) *T08H10.1 (exc-15);* (E) *C03G6.5 (exc-17);* (F) *cyk-1.* Boxed areas enlarged below each panel show areas along the canals where cytoplasm surface is swollen with vesicles, and basal surface is irregular and noticeably wider than in wild-type animals. Arrows show enlarged vesicles or cysts. Bars, 10 μm.

Knockdown of several other genes gave rise to vesicles of varying size in the cytoplasm plus irregular swellings to the side of the canal, some primarily at the terminus of the lumen, and in some cases along the length of the canals (Fig. 5B-5F). These included some animals knocked down in the F-Box gene *fbxa-183*, discussed above. Knockdowns of T08H10.1 (which will be referred to as *exc-15*) or of H09G03.1 (*exc-16*) showed increasing amounts of variable-sized vesicles in the canal cytoplasm towards the distal ends of the canals, together with increasing numbers of irregular cysts in the lumen (Fig. 5C, 5D). H09G03.1 has no conserved domains, and no clear homology to genes outside the *Caenorhabditis* genus. T08H10.1, however, encodes a well-conserved aldo-keto-reductase (family 1 member B10), and in a previous RNAi screen knockdowns of this gene slowed the defecation rate by about 20%, possibly through effects on mitochondrial stress (Liu *et al.* 2012).

Knockdown of two other genes caused the appearance of large cysts or vesicles appearing on the basal surface of the canals in just a few seemingly random spots along the length of the canals (Fig. 5E, 5F). C03G6.5 (*exc-17*) encodes another protein found only in nematodes, with a Domain of Unknown Function (DUF19), possibly an extracellular domain, found only among several nematode and bacterial proteins. CYK-1, however, is a formin of the Diaphanous class, that has a well-investigated role in regulating microfilaments in cytokinesis (Severson *et al.* 2002), and in forming normal canal morphology through interactions with the EXC-6 formin, via regulation by EXC-5 (human FGD) guanine exchange factor and CDC-42 (Shaye and Greenwald 2016). Our RNAi knockdown of *cyk-1* produced a stronger phenotype (shorter canals with large cysts on the basal side) than seen in the temperature-sensitive mutant used by the Greenwald laboratory, but not as strong an effect as was seen in double mutants of *cyk-1* (*ts*) with *exc-6* null mutants (Shaye and Greenwald 2016).

### Variability and Range of Phenotypes

Treatment of nematodes via feeding RNAi creates variable levels of knockdown between animals (Hull and Timmons 2004). This feature of the gene knockdowns has allowed observation of effects of genes that have a lethal null phenotype, and show a relationship between the phenotypes described above, as seen from RNAi-knockdown of the *vha-5* gene. This gene encodes a protein of the membrane-bound V0 subunit of the vacuolar ATPase, and is strongly expressed in the canalicular vesicles at the apical membrane of the canals (Kolotuev *et al.* 2013). Mutations of this gene are lethal, and a point mutation led to strong whorls of labelled VHA-5 at the apical surface (Liegeois *et al.* 2006). Our knockdown of this gene gave a wide range of canal phenotypes in different animals (Fig. 6). Some animals exhibited beads surrounding a normal-diameter lumen (Fig.6A), similar to animals under slow growth or osmotic stress, as in Fig. 3. Other animals showed small septate cysts in the canal lumen, but the canal lumen overall was generally of near-normal diameter, and the basal surface had mostly minor irregularities (Fig. 6B), similar to animals knocked down for *exc-15* (Fig. 5D). Other *vha-5* knockdown animals also exhibited a similar luminal phenotype, but also showed large vesicles within a highly irregularly shaped cytoplasm (Fig. 6C), similar to animals impaired in *exc-17* or *cyk-1* expression (Fig. 5E, 5F). Finally, the most extremely affected *vha-5* knockdown animals (Fig. 6D) showed cysts throughout the lumen, a swollen terminus to the lumen, and a range of variable-sized vesicles or cysts that pack the entire swollen cytoplasm of the canals. The wide range of defects seen in animals knocked down may reflect the very strong phenotype of the null mutant (embryonic lethal), and the wide range of expression of dsRNA that can occur through feeding RNAi. The variability of phenotype also indicates that the range of mutant phenotypes described above for various gene knockdowns may represent variable expression of dsRNAs that affect a common set of coordinated pathways that function to create and maintain the complicated shape of the excretory canals; these pathways include gene transcription, ion and small molecule transport, cell cytoskeleton, cell-cell communication, and movement and function of vesicles.

**Figure 6.**
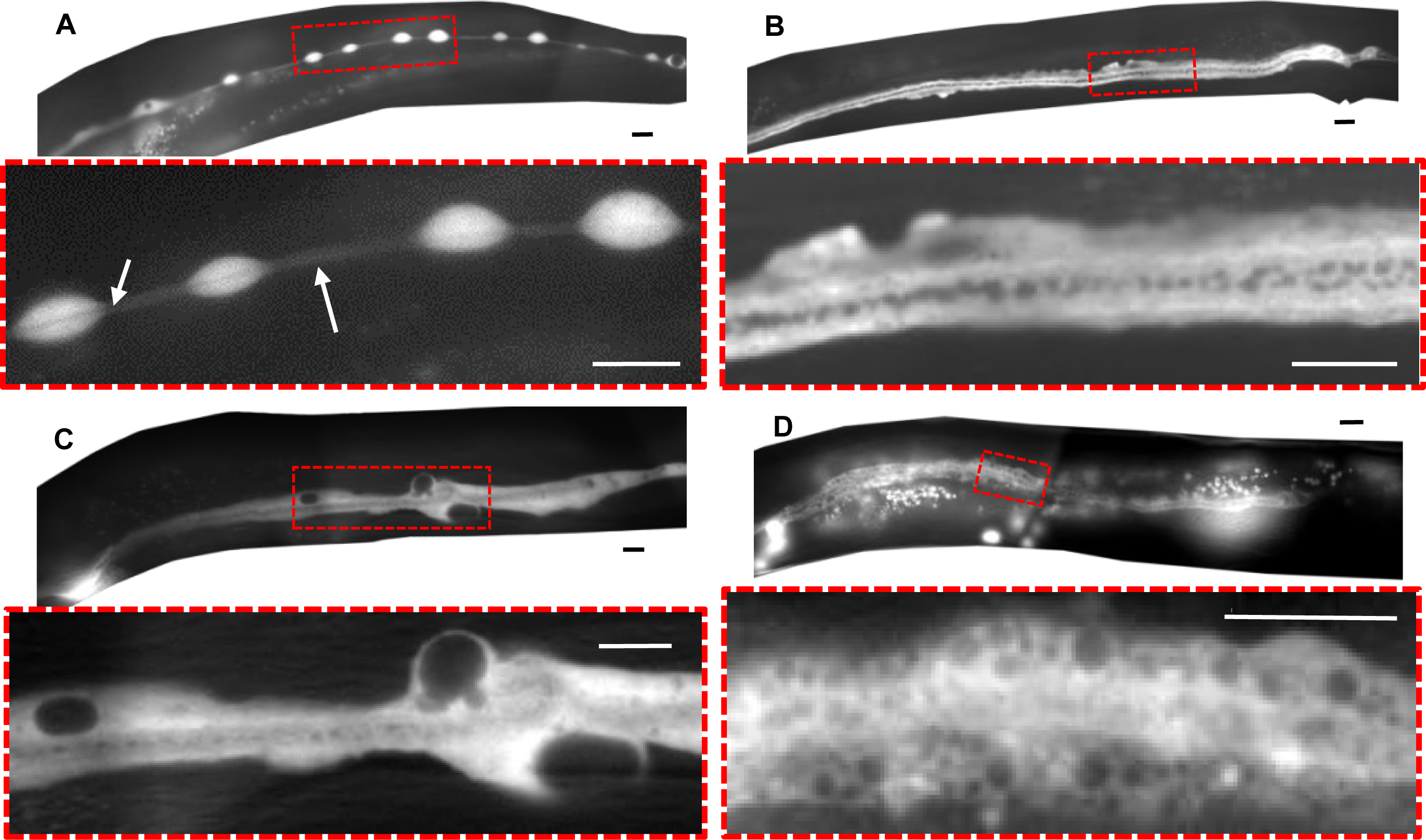
Knockdown of *vha-5* leads to a wide range of phenotypes. (A-D) GFP fluorescence of four different worms exhibiting a range of excretory canal phenotypic severity in response to *vha-5* knockdown. For each animal, the boxed area is enlarged below). (A) Periodic cytoplasmic swellings along lumen of canal. Arrows show visible lumen of normal diameter. (B) Small septate cysts in the lumen of the canal, surrounded by area of bright GFP fluorescence, and somewhat irregular diameter cytoplasm. (C) Lumen with septate cysts similar to 4B and surrounded by cytoplasm of more irregular diameter containing large cysts/vesicles. (D) Wider-diameter lumen with larger cysts, surrounded by cytoplasm filled with vesicles in a wide range of sizes. Bar, 10μm.

### Other Phenotypes

While the focus of this RNAi screen centered on excretory canal morphology, a few other phenotypes were noticed, including occasional effects on gonadal shape, fertility, and viability. In many *exc* mutants, the shape of the hermaphrodite tail spike is affected (Buechner *et al.* 1999), and similar strong results were reproducibly observed here for multiple RNAi knockdowns (Fig. 7). In addition to the knockdowns shown (for genes *exc-11, exc-14, egal-1, mop-25.2*, and *inx-12*), tail spike defects were also seen in animals knocked down in genes encoding homeobox protein CEH-6, vacuolar ATPase component VHA-5, sedoheptulose kinase EXC-10, aldo-keto reductase EXC-15, and innexin INX-13. The tail spike is formed from the interaction of hypodermal tissue hyp10 with a syncytium of two other hypodermal cells that later undergo cell death (Sulston *et al.* 1983); it remains to be determined what features this structure has in common with the canals that require the same proteins.

**Figure 7.**
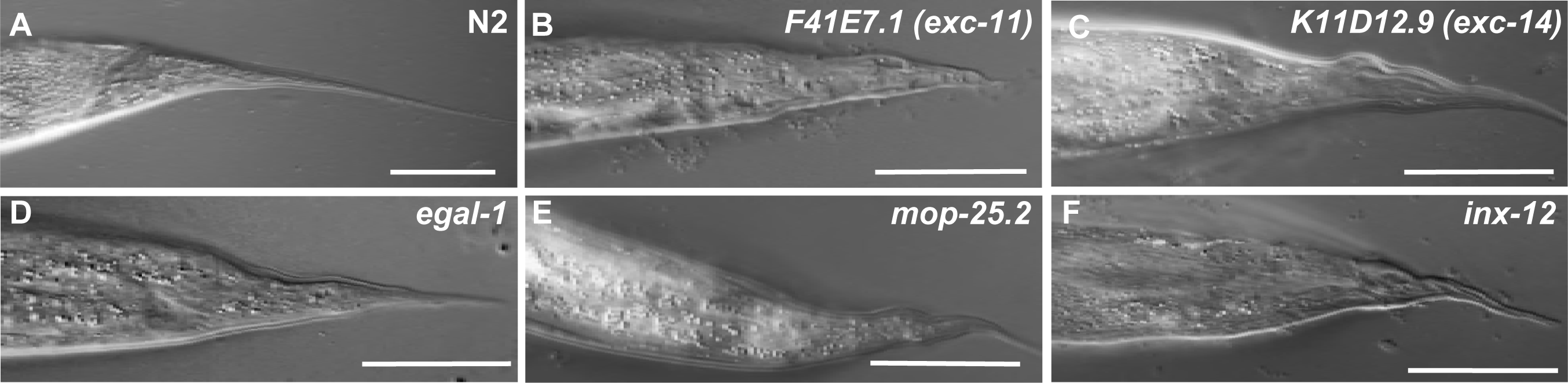
Knockdown of some *exc* genes causes tailspike defect. DIC images of the narrow tail spike of adult hermaphrodite wild-type animal (A) and of adult mutants exhibiting RNAi knockdown for genes: (B) *F41E7.1* (*exc-11*); (d) *K11D12.9* (*exc-14*); (D) *egal-1*; (E) *mop-25.2*; (F) *inx-12.* Bars, 50μm.

### Suppressors of the Exc-5 Phenotype

Finally, the RNAi screen was also carried out in animals carrying mutations in various *exc* genes, to try to find genes that interacted to form more severe phenotypes. Previous interactions have found, for example, that *exc-3; exc-7* double mutants have a more severe canal phenotype than does either mutant alone (Buechner *et al.* 1999), and similar exacerbation of effects are seen for *exc-5; exc-6* double mutants (Liegeois *et al.* 2006). No such effects were detected in this screen, but surprisingly, knockdown of two genes caused an unexpected phenotype: the restoration of near-wild-type phenotype from strongly cystic homozygous *exc-5*(*rh232*) animals (Fig. 8) carrying a large deletion of almost all of the *exc-5* gene (Suzuki *et al.* 2001). As noted above, *exc-5* encodes a Guanine Exchange Factor (GEF) specific for CDC-42 (Shaye and Greenwald 2016). EXC-5 is homologous to four human FGD proteins, including two that are implicated in Aarskog-Scott Syndrome (Facio-Genital Dysplasia) and Charcot-Tooth-Marie Syndrome Type 4H, respectively (Gao *et al.* 2001; Delague *et al.* 2007; Horn *et al.* 2012). The latter disease affects outgrowth of the single-celled tubular Schwann cells during rapid growth, and identification of mutations in suppressor genes therefore has the potential to increase understanding of this disease.

**Figure 8.**
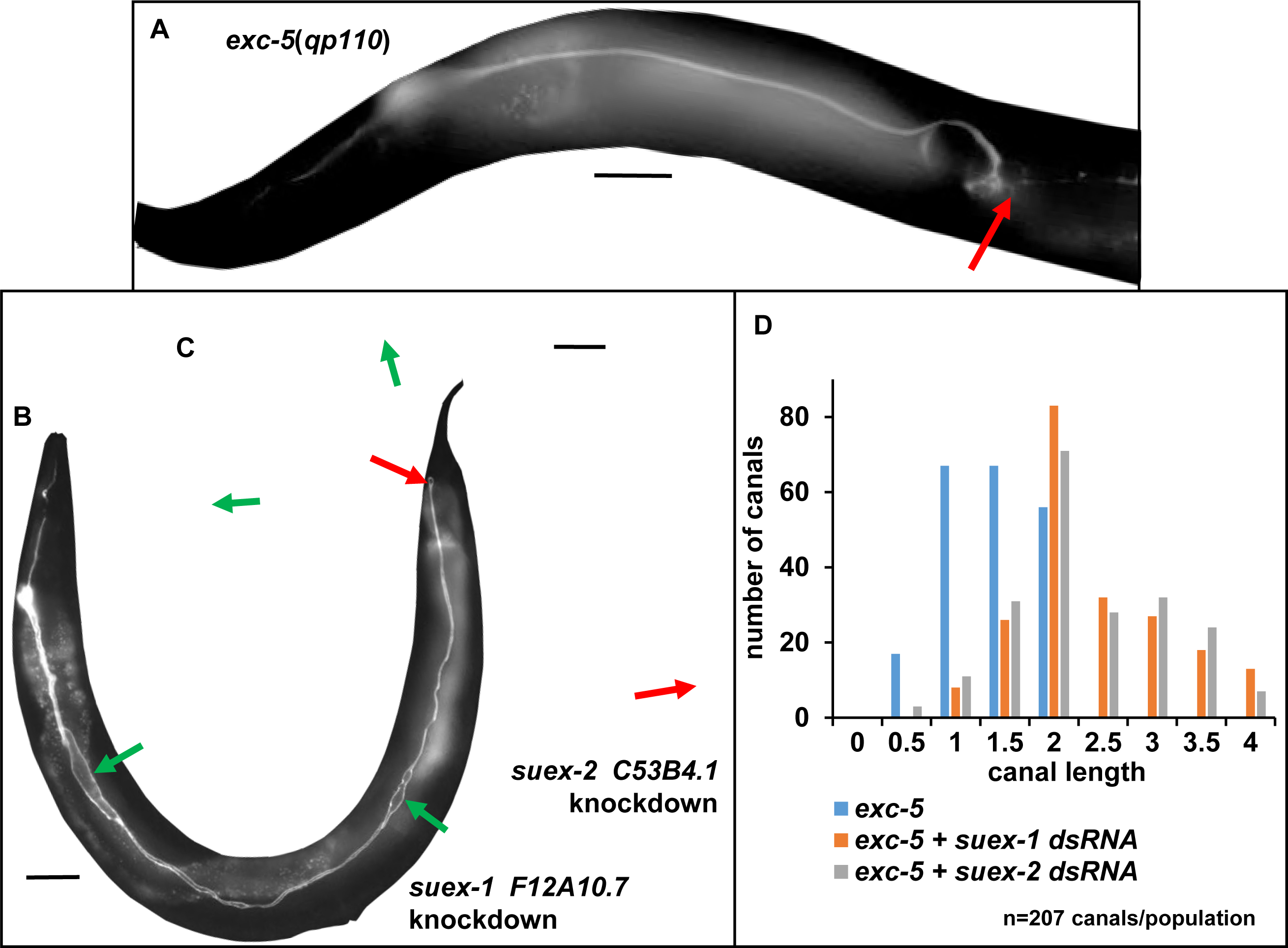
Knockdown of two genes suppresses the Exc-5 phenotype. (A-C) GFP fluorescence of canals in BK545 (null *exc-5*(*rh232*) mutants with RNAi-sensitized background and GFP expressed in canal cytoplasm (A) and of BK545 animals showing strong suppression when knocked down for (B) F12A10.7 (*suex-1*), or (C) C53B4.1 (*suex-2*). Exc-5 phenotype includes very short normal-diameter canals terminating in large cysts. Red arrows indicate termination of canals. Green arrows indicate areas of slight swelling of Suex canal lumen in both knockdowns. Bars, 50 μ,m. (D) Measurement of effect of *suex* suppression via feeding RNAi on canal length. Canals from *exc-5* mutant and mutants with *suex* knockdown were measured according to scale in Fig. 1A. Average canal length: *exc-5*(*rh232*): 1.4, *exc-5*(*rh232*); *suex-1*(*RNAi*): 2.3, *exc-5*(*rh232*); *suex-2*(*RNAi*): 2.3. N=207 for each genotype. Analysis via 3×2 Fisher’s 3×2 Exact Test (see Materials and Methods) show differences from wild-type canal length that are highly significant: P of 9.0×10^−17^ for *suex-1*, 1.7×10^−13^ for *suex-2.*

*exc-5* null mutants are characterized by large fluid-filled cysts at the terminus of both anterior and posterior canals (Fig. 8). Knockdown RNAi of these suppressor genes, both by feeding and by direct dsRNA microinjection, yielded a large number of progeny exhibiting near normal canal phenotypes, with canal length extending near-full-length (Fig. 8D). We will refer to this phenotype as Suex, for SUppressor of EXcretory defects. In SUEX canals, no obvious septate cysts are evident, although parts of the canal lumen were slightly widened (Fig. 8B, 8C).

F12A10.7 (*suex-1*) encodes a small protein (113 amino acids) unique to *C. elegans*, expressed in the excretory canal cell and in some neural subtypes, with homology to genes in only a few other *Caenorhabditis* species. The C-terminal half of the protein contains a number of repeats of tri-and tetra-peptides GGY and GGGY. As the bacterial construct from the Ahringer Library also included a small number of base pairs of the nearby gene F12A10.1, we confirmed the identity of the suppressing gene through microinjection of synthesized dsRNA specific to the F12A10.7 transcript into the gonad of the *exc-5* null mutant strain BK545, and confirmed the appearance of progeny with canals of wild-type length.

In contrast to F12A10.7, C53B4.1 (*suex-2*) encodes a protein with homologues in a wide range of animals, including humans. This gene encodes a cation transporter that has been implicated in gonadal distal tip cell migration in a previous RNAi screen (Cram *et al.* 2006). Multiple strong homologues in humans fall into the SLC (SoLute Carrier) family 22 class of proteins, with the closest homologue SLC22A1 encoding a 12-tm-domain integral membrane protein transporting organic cations (Nigam 2018) and expressed in the human liver and small intestine. The effect of knocking down this transporter implies that ionic milieu or lipid composition affects transport of vesicles mediated by EXC-5 signaling, but future work will be needed to determine the role that this transporter exerts on ionic content, and possibly on endosomal recycling in the developing excretory canal cell.

## CONCLUSION

This RNAi screen was successful at identifying 24 genes (17 not implicated before) needed to form a normal lumen of the long excretory canals of *C. elegans*. These genes encode transcription and translation factors, innexins and other channels, and proteins involved in trafficking, among others. While these processes have been implicated previously in canal tubulogenesis, these proteins identify new actors that could provide insights into how these cellular processes are integrated in single-cell tubulogenesis. Several other proteins have roles in sugar metabolism and redox, which are new processes to be involved in canal morphogenesis. Finally, two genes were identified as suppressors of *exc-5* mutation; determining the function of these suppressor proteins has the potential to increase understanding of the function of FGD protein function in normal development and in disease.

## ACKNOWLEDGMENTS

H.A. was supported in part by KU Graduate Research Funds #2301847 and #2144091. E.A.L. was supported by National Institutes of Health grants #NS0090945, NS0095682, NS0076063, and GM103638. Some strains were provided by the CGC, which is funded by the NIH Office of Research Infrastructure Programs (P40 OD010440). RNAi-refractive strain NL3321 *sid-1*(*pk3321*) was the gift of Lisa Timmons, U. Kansas. Some strains were created by the *C. elegans* Reverse Genetics Core Facility at the University of British Columbia, which is part of the international *C. elegans* Gene Knockout Consortium.

## AUTHOR CONTRIBUTIONS

M.B. and H.A. designed the research, with support from E.A.L. on strategies for RNAi treatments; H.A. and T.C. performed bacterial and nematode growth, performed RNAi treatments, and evaluation of canal phenotypes. E.A.L. supplied bacterial RNAi clones. M.B. and H.A. prepared the figures and wrote the manuscript.

